# Bridging Big Data: Procedures for Combining Non-equivalent Cognitive Measures from the ENIGMA Consortium

**DOI:** 10.1101/2023.01.16.524331

**Authors:** Eamonn Kennedy, Shashank Vadlamani, Hannah M Lindsey, Pui-Wa Lei, Mary Jo-Pugh, Maheen Adamson, Martin Alda, Silvia Alonso-Lana, Sonia Ambrogi, Tim J Anderson, Celso Arango, Robert F Asarnow, Mihai Avram, Rosa Ayesa-Arriola, Talin Babikian, Nerisa Banaj, Laura J Bird, Stefan Borgwardt, Amy Brodtmann, Katharina Brosch, Karen Caeyenberghs, Vince D Calhoun, Nancy D Chiaravalloti, David X Cifu, Benedicto Crespo-Facorro, John C Dalrymple-Alford, Kristen Dams-O’Connor, Udo Dannlowski, David Darby, Nicholas Davenport, John DeLuca, Covadonga M Diaz-Caneja, Seth G Disner, Ekaterina Dobryakova, Stefan Ehrlich, Carrie Esopenko, Fabio Ferrarelli, Lea E Frank, Carol Franz, Paola Fuentes-Claramonte, Helen Genova, Christopher C Giza, Janik Goltermann, Dominik Grotegerd, Marius Gruber, Alfonso Gutierrez-Zotes, Minji Ha, Jan Haavik, Charles Hinkin, Kristen R Hoskinson, Daniela Hubl, Andrei Irimia, Andreas Jansen, Michael Kaess, Xiaojian Kang, Kimbra Kenney, Barbora Keřková, Mohamed Salah Khlif, Minah Kim, Jochen Kindler, Tilo Kircher, Karolina Knížková, Knut K Kolskår, Denise Krch, William S Kremen, Taylor Kuhn, Veena Kumari, Jun Soo Kwon, Roberto Langella, Sarah Laskowitz, Jungha Lee, Jean Lengenfelder, Spencer W Liebel, Victoria Liou-Johnson, Sara M Lippa, Marianne Løvstad, Astri Lundervold, Cassandra Marotta, Craig A Marquardt, Paulo Mattos, Ahmad Mayeli, Carrie R McDonald, Susanne Meinert, Tracy R Melzer, Jessica Merchán-Naranjo, Chantal Michel, Rajendra A Morey, Benson Mwangi, Daniel J Myall, Igor Nenadić, Mary R Newsome, Abraham Nunes, Terence O’Brien, Viola Oertel, John Ollinger, Alexander Olsen, Victor Ortiz García de la Foz, Mustafa Ozmen, Heath Pardoe, Marise Parent, Fabrizio Piras, Federica Piras, Edith Pomarol-Clotet, Jonathan Repple, Geneviève Richard, Jonathan Rodriguez, Mabel Rodriguez, Kelly Rootes-Murdy, Jared Rowland, Nicholas P Ryan, Raymond Salvador, Anne-Marthe Sanders, Andre Schmidt, Jair C Soares, Gianfranco Spalleta, Filip Španiel, Alena Stasenko, Frederike Stein, Benjamin Straube, April Thames, Florian Thomas-Odenthal, Sophia I Thomopoulos, Erin Tone, Ivan Torres, Maya Troyanskaya, Jessica A Turner, Kristine M Ulrichsen, Guillermo Umpierrez, Elisabet Vilella, Lucy Vivash, William C Walker, Emilio Werden, Lars T Westlye, Krista Wild, Adrian Wroblewski, Mon-Ju Wu, Glenn R Wylie, Lakshmi N Yatham, Giovana B Zunta-Soares, Paul M Thompson, David F Tate, Frank G Hillary, Emily L Dennis, Elisabeth A Wilde

## Abstract

Investigators in neuroscience have turned to Big Data to address replication and reliability issues by increasing sample sizes, statistical power, and representativeness of data. These efforts unveil new questions about integrating data arising from distinct sources and instruments. We focus on the most frequently assessed cognitive domain - memory testing - and demonstrate a process for reliable data harmonization across three common measures. We aggregated global raw data from 53 studies totaling N = 10,505 individuals. A mega-analysis was conducted using empirical bayes harmonization to remove site effects, followed by linear models adjusting for common covariates. A continuous item response theory (IRT) model estimated each individual’s latent verbal learning ability while accounting for item difficulties. Harmonization significantly reduced inter-site variance while preserving covariate effects, and our conversion tool is freely available online. This demonstrates that large-scale data sharing and harmonization initiatives can address reproducibility and integration challenges across the behavioral sciences.

**Teaser:** We present a global effort to devise harmonization procedures necessary to meaningfully leverage big data.

## Introduction

Data sharing consortia aim to increase the robustness and statistical power of results by aggregating large and diverse samples.(*1, 2*) While analyses of large datasets can provide unparalleled statistical power, data aggregation without robust harmonization can mask and even introduce flaws and biases.(*3*) This is a critical consideration in the behavioral sciences, where multi-site collaboration often requires the synthesis of non-identical cognitive measures.(*4*–*6*) For example, verbal memory/recall is a core cognitive function, and deficits in learning and memory are one of the most common and widely assessed patient complaints.(*7*) However, a wide variety of auditory verbal learning tasks (AVLTs) exist that can be administered to assess verbal memory and recall, and these differ across a range of qualitative and quantitative features.(*7*–*9*) Such differences in assessment instruments can contribute to inconsistencies in the measurement of neurocognitive performance.(*10*)

Methods to accurately convert scores across AVLTs could facilitate highly powered studies of verbal memory and recall, offering opportunities for new clinical insights.(*11, 12*) However, data from single sites are typically biased by the specific attributes, demographics, and inclusion criteria of the study which can increase variance/error and confound reproducibility.(*13–15*) To address these limitations, emerging data harmonization approaches offer new ways to perform data transformations that remove unwanted influences in aggregated data while preserving meaning. Data harmonization of large and heterogeneous AVLT data sources represents an appropriate framework for the development of cross-AVLT score conversion tools, but such efforts come with analytical and organizational challenges.(*1, 4, 16, 17*)

Here, we report a retrospective multisite (n = 53 datasets) mega study analysis of three Common AVLTs: the California Verbal Learning Test (CVLT),(*7*) the Rey Auditory Verbal Learning Test (RAVLT),(*18*) and the Hopkins Verbal Learning Test-Revised (HVLT),(*19*) drawing from international healthy and brain-injured populations across 13 countries and 8 languages. In contrast to meta-analyses which combine summary statistics from several sites, mega-analysis centralizes and pools individual raw data from many sites prior to analysis. This allows for a richer range of experimental designs which can consider subtle single item differences in detail.(*1, 2*) Our primary hypothesis was that conversion performance would be significantly improved by a mega-analytic pipeline combining harmonization and item response theory (IRT) relative to unadjusted conversions. The goal of this study was to establish crosswalks between common memory measures as a means to address long standing data compatibility issues for AVLTs through freely available conversion tools: https://verbal-learning.halfpipe.group/

## Results

### Data Summary

An overview of the key features differentiating the CVLT, RAVLT, and HVLT assessments are provided in Table 1. Supplementary Table S1 shows summary characteristics itemized for each of the 53 aggregated datasets after applying exclusion criteria. Overall, the sample size was N = 10,505 (31.8% female) which included both controls and TBI groups. The median age was 42 years with an interquartile range of 30-55 years. Studies showed significant differences in both item scores and covariates (Fig. 1). Table 2 shows the summary statistics of the full cohort after aggregation. Each instrument was represented by >1000 subjects across >10 studies, indicating good representation of AVLTs. Significant differences in demographic characteristics were evident across measures, indicating that covariate adjustment was required.

**Figure 1.**
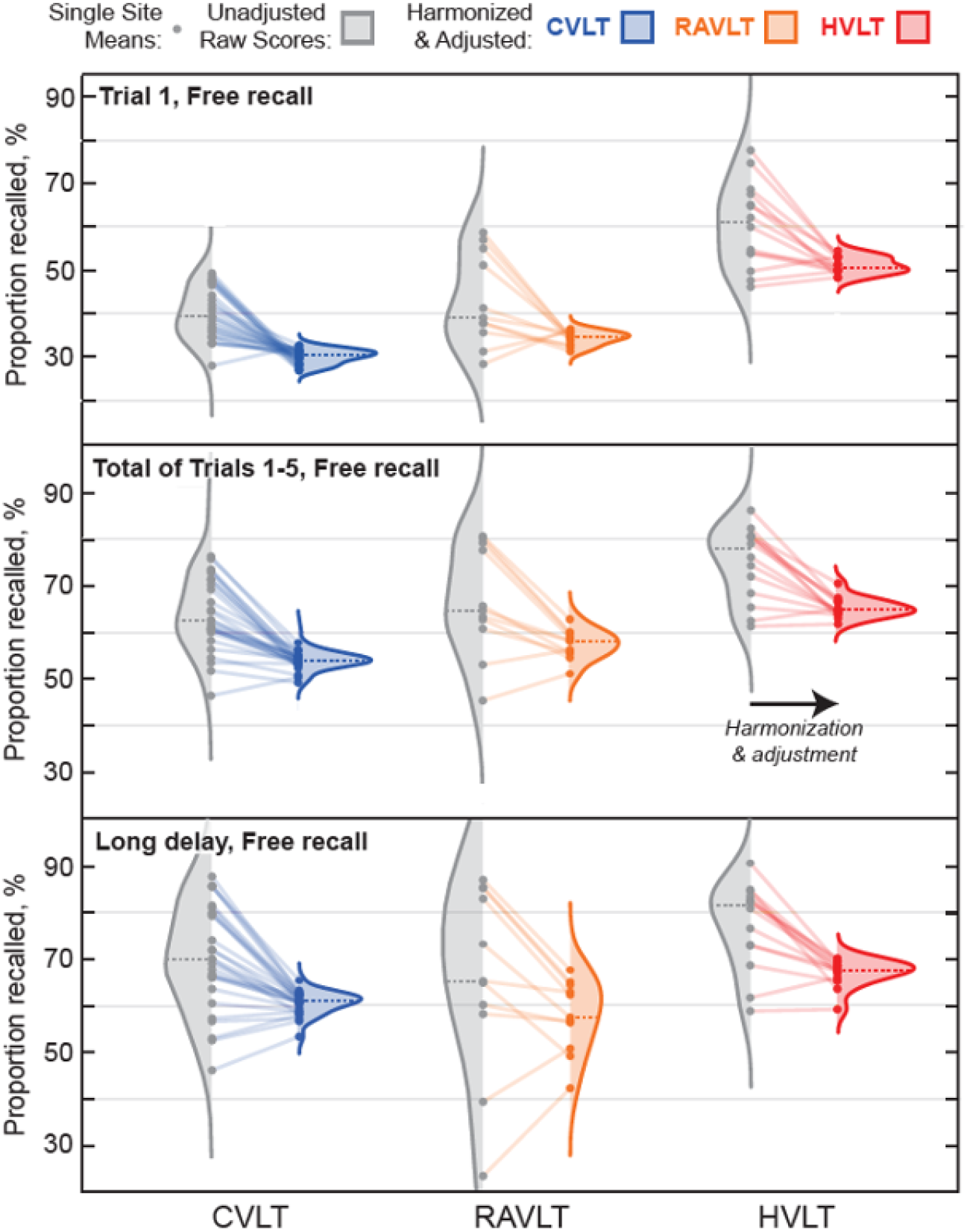
Comparing proportions of memory items recalled before and after harmonization. Mean scores for each site (dots) are shown broken out by instrument (color) and item (Top: Trial 1 immediate free recall, Middle: Total sum of all Trials, Bottom: Long-delay free recall scores).

**Table 1.** Summary of the key features of the three AVLTs.

|  | CVLT | RAVLT | HVLT |
| --- | --- | --- | --- |
| List lengths (words) | 16 | 15 | 12 |
| Semantic categories | 4 | - | 3 |
| Immediate free recall trials | 5 | 5 | 3 |
| Recognition trial, total words | 48 | 50 | 24 |
| Long delay duration (minutes) | 20-30 | 20 | 20-25 |
| Distractor list (List B) | Yes | Yes | No |
| Cued recall trials | Yes | No | No |
| Short Delay Trial | Yes | Yes | No |

### Harmonization

The ComBat-GAM algorithm was implemented to correct for site-specific variations such as differences in inclusion/exclusion criteria (Supplementary Table S2) while explicitly preserving real covariate effects (e.g., age) for further analysis as described elsewhere.(*4*) Fig. 1 shows a comparison of single site mean scores, before (gray dots) and after (colored dots) site harmonization. A line is drawn to connect each site from its pre- to post-harmonized mean value. Gray distributions portray the variation in mean scores across sites before harmonization; colored distributions portray site mean score distributions after harmonization (CVLT: Blue, RAVLT: Orange, HVLT: Red). The unadjusted distributions of scores (gray areas) exhibit much higher variance than their post-harmonized equivalents, and overall harmonization reduced total variance by 37% across all items.

### Covariate Adjustment

Fig. 2a shows boxplots of sum of raw learning trial scores as percentages stratified by group (TBI vs. control) and sex/gender. TBI history and male sex/gender were both associated with lower sum scores across all three tests. Age-related declines were well-fit by quadratics; years of education were well-fit by a straight line. The effects of both age and education were significant and consistent across all tests. Linear models within each measure were used to assess covariates (Table 3) after language and country of origin effects were removed separately prior to harmonization.

**Figure 2.**
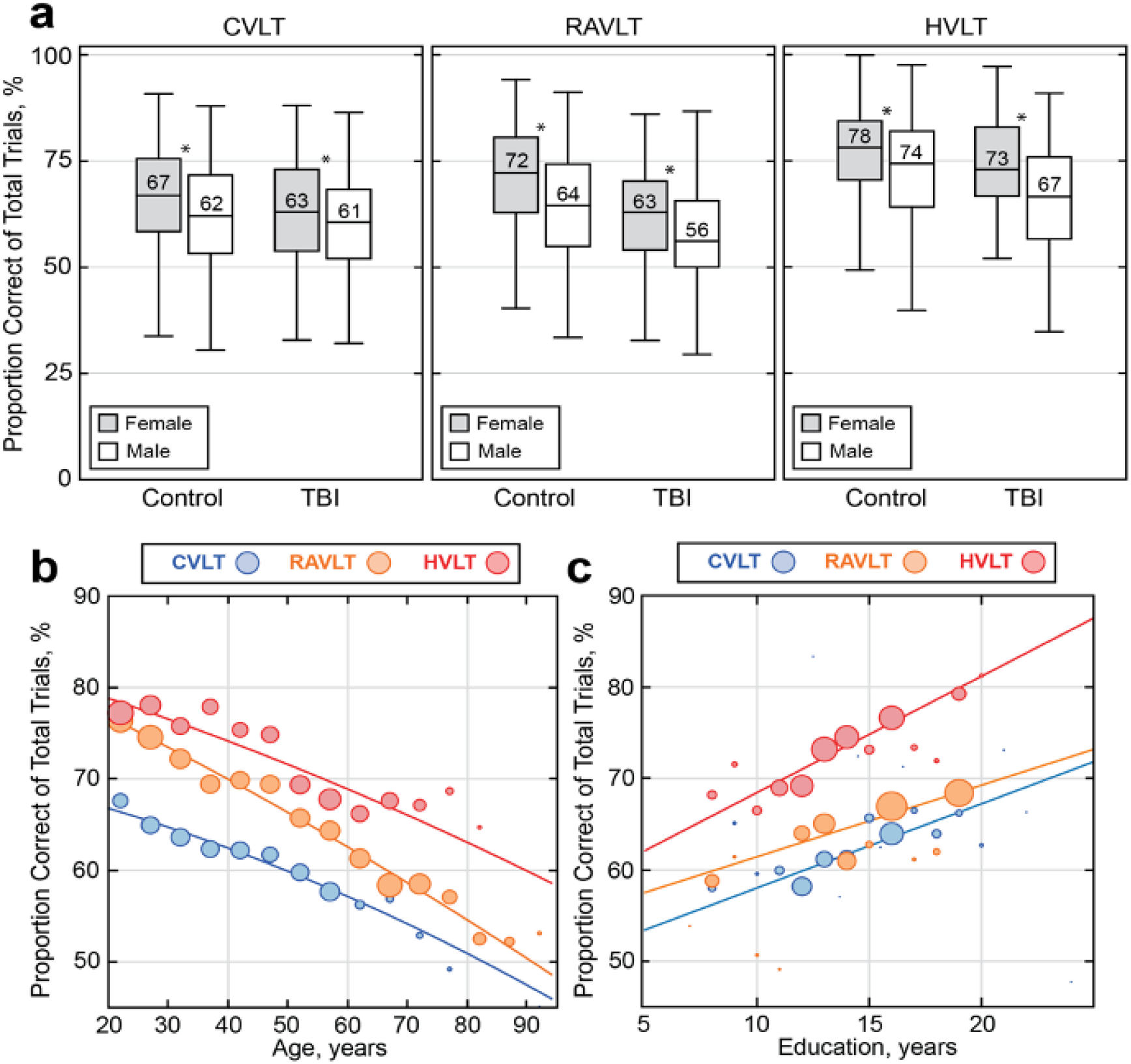
Visualizing covariate effects on scores across AVLTs. **(a)** Boxplots of scores stratified by group (TBI vs. control) and sex/gender indicated that males and those with history of TBI had significantly lower scores on average for all tests. Age-related declines **(b)** and the beneficial effects of education **(c)** on scores were consistent across all AVLTs.

**Table 2.** Descriptive characteristics of the total cohort by instrument.

|  | **CVLT** | **RAVLT** | **HVLT** | ***p*** |
| --- | --- | --- | --- | --- |
| Sample size, N | 6,634 | 2,728 | 1,143 |  |
| # Datasets | 28 | 11 | 14 |  |
| Age: 16-35 | 34.70% | 32.70% | 48.20% | <0.001 |
| 35-65 | 61.20% | 40.40% | 41.90% | <0.001 |
| 65+ | 4.10% | 26.90% | 9.90% | <0.001 |
| Gender: Male | 76.50% | 48.60% | 66.10% | <0.001 |
| Female | 23.50% | 51.40% | 33.90% | <0.001 |
| Region: Americas | 90.70% | 44.50% | 69.00% | <0.001 |
| Europe | 8.30% | 55.40% | 27.60% | <0.001 |
| Asia Pacific | 1.10% | 0.10% | 3.40% | <0.001 |
| Language: English | 90.70% | 37.60% | 82.20% | <0.001 |
| Spanish | 1.90% | 7.40% | 6.40% | <0.001 |
| German | 0.20% | 35.3% | 11.50% | <0.001 |
| Italian | 0.00% | 8.80% | 0.00% | <0.001 |
| Other | 7.20% | 10.90% | 0.00% | <0.001 |
| Education: Secondary | 20.80% | 28.30% | 15.20% | <0.001 |
| Some College | 14.80% | 14.00% | 19.70% | <0.001 |
| Postgraduate | 7.20% | 21.40% | 5.50% | <0.001 |
| Clinical Population: TBI | 64.70% | 11.90% | 21.90% | <0.001 |
| Controls | 35.30% | 88.10% | 78.10% | <0.001 |
| Mean Raw Scores: Trial 1 | 4.9 ± 1.9 | 5.0 ± 1.9 | 6.1 ± 1.7 | <0.001 |
| Total Learning | 43.2 ± 9.7 | 42.9 ± 8.6 | 23.2 ± 4.2 | <0.001 |
| Long Delay | 9.6 ± 3.1 | 8.7 ± 3.0 | 7.9 ± 2.2 | <0.001 |

**Table 3.** Blocked linear regressions predicting sum of raw learning scores per instrument. The average percentage effect across all AVLTs are shown, indicating that Age>65 had the largest impact on scores overall (−11.4%). * indicates significance after correction for multiple comparisons.

|  | CVLT | RAVLT | HVLT |  |  |
| --- | --- | --- | --- | --- | --- |
| Sample size | 6,634 | 2,728 | 1,143 |  |  |
| Explained variance | 10% | 30% | 22% | % effect | Significant |
| Max score of T1-TN | 80 | 75 | 36 | all | all tests |
| **Age group (Ref: 16-35 years)** |  |  |  |  |  |
| 35-65 | -2.7* | -6.5* | -1.0* | -4.90% | Yes |
| 65+ | -7.8* | -12.2* | -2.9* | -11.40% | Yes |
| **Sex/gender (Ref: Male)** |  |  |  |  |  |
| Female | 3.1* | 3.7* | 1.6* | 4.40% | Yes |
| **Race/Ethnicity (Ref: White)** |  |  |  |  |  |
| Multi | -1.3 | -3 | -0.6 | -2.40% | No |
| Black | -2.7* | -0.8 | -2.8* | -4.10% | No |
| Asian | 1.1 | 0.5 | -1.9* | -1.10% | No |
| Hispanic | 0.5 | -1.1 | -0.1 | -0.40% | No |
| **TBI (Ref: No TBI)** | -1.3* | -1.6* | -1.6* | -2.80% | Yes |
| **Education (per year)** | 0.6* | 0.6* | 0.4* | 0.90% | Yes |

### Anchor Items and Ability Measures

Anchor items similar in format and nature were identified for each of the three required crosswalks (1. RAVLT ↔ CVLT; 2. RAVLT ↔ HVLT; and 3. CVLT ↔ HVLT). After expert consensus and trials of different anchor combinations, we elected to use immediate free recall learning trials, short delay, and long delay free recall as anchor items, where available. Short delay was used as an anchor item between CVLT and RAVLT only (short delay is not assessed in HVLT). False positive measures were not recorded consistently across sites and were not used. Recognition hits showed inconsistent behavior and were excluded from conversions (see Limitations). Since all site effects and measured covariate effects had been removed prior to IRT analysis, we assumed scores were distributed randomly across measures and ability scores did not require further scaling.

### Item Response Theory

A continuous IRT model (Fig. 3) was used to estimate the latent trait of all individuals while accounting for different item difficulties and discriminations across tests. Statistically, individuals with the same verbal learning ability will achieve equivalent scores on different measures. Third degree polynomials estimated the relationship between observed scores and ability scores for each. Cubic polynomial fits of ability vs. score are shown in Fig. 3a for immediate, short, and long delay items. Horizontal lines of equivalent ability connect equivalent scores. Items with longer delay show larger differences in difficulty across AVLTs.

**Figure 3.**
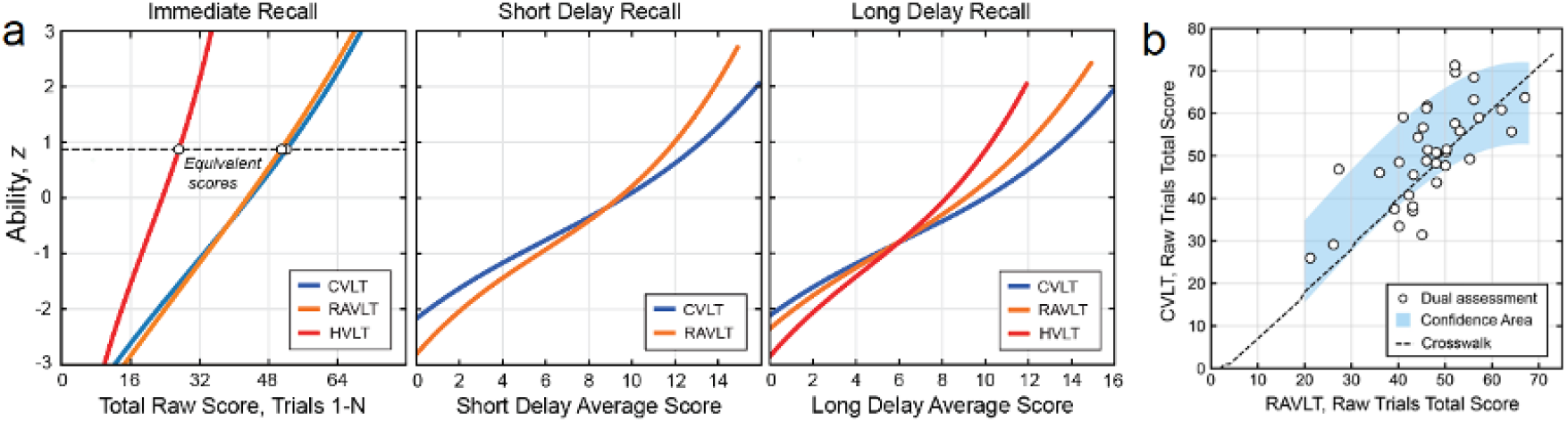
Visualizing and Validating Conversions. **(a)** Average raw scores as a function of individual ability are shown approximated as cubic polynomial fits for immediate, short, and long delay trials. Horizontal lines of equivalent ability connect equivalent raw scores across tests, which facilitates the construction of crosswalks. **(b)** Scatter plot and fit of the adjusted raw sum of learning Trial scores for a subset of cases who were administered both the CVLT and RAVLT (n=36). The confidence area of the dually assessed data is shown in blue, and agrees with the derived crosswalk for CVLT->RAVLT (n=9,362, black dotted line).

### Validation with Dually Administered Tests

We validated the derived conversions on held-out data not used in other analyses. Validation was conducted by comparing the conversion estimates to real data where two verbal learning tests were administered to the same set of individuals (Fig. 3b; n=36). How well conversion lines fit the dually administered test scores is an independent measure of conversion accuracy.

Fig. 3b shows a scatter plot of the raw sum of learning scores for a subset of cases who were administered both the CVLT and RAVLT after harmonization. As a proportion correct, repeated assessment scores vary within subjects by 10.7% on average, presented as a confidence area derived by 100x cross validation assuming a fit through the origin (Fig. 3b, blue shaded area). These data are fitted against the IRT-derived conversion scores (black dotted line) for RAVLT to CVLT. The line falls within the 95% confidence bound for the dually administered tests, indicating agreement. Compared to the same conversion model constructed using unadjusted data, the harmonized conversion exhibited a 9.5% lower root mean squared error against the held-out data, indicating that harmonization moderately improved conversion.

### Potential Clinical Application

An individual with a chief complaint of memory loss is undergoing evaluation to address a question of possible neurodegenerative disease. Initial neuropsychological examination revealed a score on the CVLT suggesting memory impairment. However, the individual is seeking a second opinion with clinicians at another institution who elect to repeat memory testing. The second neuropsychologist administers the RAVLT. Using the online conversion tool the RAVLT is converted to an equivalent CVLT score, and these scores can now be directly compared after the same CVLT norms are applied.

## Discussion

There have never been more studies published annually in the history of the neurosciences.(*20*) This intensive rate of research offers unparalleled opportunity for data combination and nuanced examination of cognitive and behavioral changes associated with neurological diagnoses. However, “high volume science” lacks coordination between studies, which poses critical challenges for the integration of findings and data harmonization. For example, AVLTs are the most common method for learning and memory assessment, but they were independently developed and without explicit quantitative reference to pre-existing instrumentation. Over the last 70 years, this has led to a scenario where clinicians and researchers routinely use distinct AVLTs with incomparable results.(*7*–*9*) This is not only a technical inconvenience but is problematic for the interpretation and reproducibility of results and findings.

Constructing reliable standards for converting scores across common AVLTs is challenging, because conversions should be made independent of factors such as language, study group, and instrumental details. For example, given more words to recall, it is more likely that more words will be recalled. Naim et al. found the average number of memory items recalled (R) scales with the root of M items presented,(*21*) and there are other subtle differences between seemingly similar assessments (Table 1). Together, large-sample mega-analysis and harmonization present a promising solution to address these concerns in aggregate, offering new opportunities to explore previously incompatible datasets and examine more interesting clinical features.

Beyond conversions, comparing the difficulty of tests on the same ability scale (Fig. 3a) may assist with the selection of AVLT across different research and clinical contexts. For example, the HVLT was the easiest test overall, while the CVLT was the most challenging test. The HVLT may be most appropriate for the assessment of individuals who are at risk for significant impairment. Conversely, the CVLT has sufficient dynamic range to discriminate within high ability groups, while the RAVLT may be well suited for studies involving a wide range of abilities. However, these are relatively coarse recommendations, and individually-derived measures from the RAVLT and other AVLTs may be more or less suitable given specific scenarios.(*22*)

In the process of converting across AVLTs, site effects such as different settings, inclusions, and procedures were found to have an appreciable impact on verbal learning scores (Fig. 1, gray distributions). This may also be attributable to underlying differences in inclusion/exclusion criteria across studies (Table S2). However, a detailed list of all the ways our sources differed was not necessary to remove these effects in aggregate with a harmonization algorithm. We confirmed our primary hypothesis that conversion error would be reduced by implementing a mega-analytic pipeline combining harmonization and IRT. In the end, simple cubic models effectively equated the instrumental component of scores without reference to underlying factors (Fig. 3a). In time, these conversions may be found to be suitable for clinical utilization at the individual level, although verifying this will require further independent scrutiny (see Limitations).

Appropriate harmonization transforms data in ways that preserve its core relationships.(*5*) For this study, these relationships include the associations between scores and ability, between scores and covariates (e.g., age-related memory decline), and the measurement of the underlying cognitive construct. These relationships can be assessed directly by comparing pre/post harmonized data (Fig. 1). After harmonization, the higher scores associated with younger age, female sex/gender, more education, and controls persisted for all AVLTs, despite a large drop in cross-site variance. Interestingly, despite the large sample size (to our knowledge, the largest AVLT dataset to date), no Race/Ethnicity variable was consistently associated with higher or lower scores (Table 3).

Beyond covariate effects, harmonizing AVLT offers opportunities for new, highly powered mega-analytic investigations of verbal learning and memory. For example, aggregating previously incompatible data sources could yield new estimates of the impact of a range of clinical conditions on the ability to encode, combine, store, and retrieve information. This may be particularly beneficial as a means to functionally characterize the imaging findings from large-scale ENIGMA mega studies, where interesting imaging features are emerging in clinical populations not seen in smaller samples (*23*) Large-scale harmonization efforts also hold the promise of advancing theory. For example, through common definitions of cognitive constructs, these efforts may influence cognitive theory by advancing the development of cognitive ontologies.(*24*)

## Limitations

Individuals who took different tests may have different abilities on average, meaning some residual correlation between ability and choice of instrument may have persisted. However, a post-hoc analysis of converted scores found that the average percentage difference in scores across AVLTs was just 0.052%.

This study considered only a limited binary interpretation of lifetime history of TBI, which exists along a spectrum of severity and has distinct phenotypes.(*25*) For example, late effects of TBI are highly variable with injury severity a major factor (e.g chronic memory/learning deficits are expected with severe injury but are uncommon with mild TBI). A secondary sensitivity analysis conducted using only the control population and no individuals with TBI resulted in similar conversions.

This study aggregated 53 international datasets, but these were primarily from English-speaking and western hemisphere countries, and this study should not be assumed to be a true internationally balanced sample. Our IRT conversions were validated against held-out data of dually administered tests (Fig. 3b) which found that the harmonization pipeline reduced conversion error by 9.5% compared to unadjusted conversions. However, we did not have data to independently assess the other two conversions (RAVLT ↔ HVLT and 3. CVLT ↔ HVLT), which limits the strength of this external check.

We attempted to construct a crosswalk for recognition memory trials, but unlike the other items, we could not establish low error IRT results for the recognition item and the recognition hits memory trial conversions were not included.

## Conclusion

To address long standing incompatibilities across commonly used AVLTs, we aggregated a comprehensive dataset of more than ten thousand participants drawn from 53 international datasets that recorded performance on verbal learning tasks. We confirmed our hypothesis, and found that site harmonization and IRT could isolate the instrumental component of scores. This work suggests the specific choice of AVLT has a pronounced effect; Averaged across items, the CVLT was the most challenging test, although it was similar to the RAVLT in difficulty, while the HVLT was the least difficult, as expected due to its lower complexity.(*26*) Free conversion tables and tools (available at verbal-learning.chpc.utah.edu) can assist clinicians to track and compare patient raw scores against large reference groups, regardless of differences in AVLT administration practices. More broadly, this work demonstrates that data harmonization of large data sharing initiatives can offer new tools and a means to address long standing data challenges across the behavioral sciences.

## Materials and Methods

### Data Sources and Inclusion Criteria

Drawing from a range of international studies of head injury and comparator groups and controls for a variety of conditions (*27–31*) this secondary multisite (N = 53 datasets) mega study analysis focuses on three AVLTs: the CVLT,(*7*) HVLT,(*19*) and RAVLT.(*18*) To mitigate balance issues, we included only comparator and groups with TBI to reduce the mean difference in ability across assessments. As described in prior work,(*4*) we aggregated data contributed by collaborators in the Psychiatric Genomics Consortium (PGC), the Enhancing NeuroImaging Genetics through Meta-Analysis Consortium (ENIGMA) working groups,(*32*) the ENIGMA Brain Injury working group,(*15*) and the Long-term Impact of Military-relevant Brain Injury Consortium – Chronic Effects of Neurotrauma Consortium, LIMBIC-CENC.(*17*) The University of Utah provided overall Institutional Review Board (IRB) study approval. Inclusion criteria, performance validity testing, and other factors varied across studies (see Supplementary Table S2).

To limit sources of variability, we excluded anyone with a known clinically diagnosed mental health or neurological condition other than traumatic brain injury (TBI). Consistent with standard AVLT administration practices, we included only participants aged 16 years or over. In the case of longitudinal or serial measurement designs, only the first measurement of AVLTs per person were included; repeated measurements were dropped.

### Verbal Learning Task Contents and Scoring

Table 1 provides a summary overview of the key features of the AVLTs assessed. AVLT scores on each trial denote the number of correct words that are recalled. The maximum score reflects the number of memory items per list. The sum of the total words recalled across all immediate free recall (learning) trials is the immediate free recall summary score (Sum of Learning Trials). These raw scores are often subsequently normed so that the performance of the individual can be contextualized relative to a population of interest. However, in this work we exclusively assess raw scores, and not t-scores, or normative scores. We focus on raw scores because normative values are occasionally updated over time and are based upon distinct normative samples between instruments.

### The California Verbal Learning Test

The CVLT (*7*) refers to a family of instruments that assess verbal learning and memory deficits. The CVLT has been revised twice, and three iterations exist (CVLT-I, CVLT-II, and CVLT-3). Additionally, the CVLT comes in standard, short, and alternate forms. In this work, we estimated crosswalks for the more recent CVLT-II and the CVLT-3. While the CVLT-3 is nominally a revision of the CVLT-II, in practice the target words, their order, and their number are the same for both the CVLT-3 and CVLT-II. Thus, we refer to both CVLT-3 and CVLT-II standard and alternate forms together as ‘CVLT’. Table 1 provides a numerical overview of the key features of the CVLT. The CVLT uses M = 16-word list lengths, which are drawn from 4 semantic categories, and 5 consecutive learning trials. The CVLT is a comprehensive test that includes a distractor list, cued and free recall assessments, short and long delay trials, and a recognition trial with 48 words.

### The Hopkins Verbal Learning Test–Revised

The HVLT-R (*19*) is a relatively short measure of verbal learning and memory deficits. The HVLT exists in two primary forms (original and revised) denoted together as ‘HVLT’. Table 1 outlines the key features of the HVLT. The HVLT does not use a distractor list for immediate recall and does not assess cued or a short delay recall performance. The HVLT uses M = 12-word list lengths, which are drawn from 3 semantic categories, and uses a small (N=24) total pool of words for scoring. The HVLT has three consecutive learning trials.

### The Rey Auditory Verbal Learning Test

The RAVLT (*18*) is a measure of verbal learning and memory deficits. Table 1 provides an overview of the key features of the RAVLT. The RAVLT draws from random, semantically unrelated words, and employs a M = 15-word list length, a distractor list, as well as a large (N=50) total pool of words for scoring recognition hits. Alternate forms also exist for the RAVLT.

### Covariates

Language, country of origin, age at testing, sex/gender, race/ethnicity, site/study, military/civilian status, TBI history, and education level were included and adjusted for in this study. The exclusion criteria were used to rule out the presence of any other clinically relevant variables, including epilepsy, dementia, and mild or early onset cognitive impairment.(*33*) While some of the studies recorded gender, others recorded biological sex, and these were aggregated into a single variable. Ethnicity was binarized to Hispanic/Latino, or Not Hispanic/Latino. Perspectives on race/ethnicity differ widely according to cultural context,(*34*) and we elected to use broad categories of Black, White, Asian, and Other. Covariate coefficients per AVLT model were converted to percentages, averaged, and then applied back to adjust the full cohort. This means each covariate had the same effect on scores regardless of the instrument used.

### Statistical Analysis

Analysis was performed in Python 3 and in R. Kruskal–Wallis *H* tests (omnibus) were used to test for overall significance across groups. Where normality was confirmed, Welch’s *t* tests were used for post-hoc pairwise comparisons with additional correction for multiple comparisons. Overall missing values were low (<5%) and any missing data points were imputed with nearest neighbor imputation. After data cleaning and imputation, Empirical Bayes harmonization using the ComBat-GAM algorithm (*13*) was used to remove unwanted site effects while preserving instrumental effects for further analysis.

### Site and Covariate Modeling

Modeling was conducted in three stages: first, the overall dataset was divided into three subsets, one per AVLT instrument. The ComBat-GAM algorithm (*13*) was applied to each of these three subsets separately to remove site effects within each AVLT. Prior work evaluating the harmonization of cognitive measurements with this approach found that harmonization preserved real covariate effects while removing site effects.(*5*) After site correction, covariate adjustment was performed as follows: the overall dataset was again divided into three distinct subsets by instrument, and ordinary least squares (OLS) linear models were used to estimate and remove covariates averaged across all three AVLT linear models.

### Item Response Theory

A Continuous Model IRT family (*23, 35–37*) was used to estimate each subject’s verbal learning ability because item score ranges were different across tests. To calibrate parameters, we used Shojima’s (*36*) simplified expectation maximization (EM) method by assuming non-informative priors for item parameters as implemented in the EstCRM (Continuous Response Model) R package.(*35*) After all data adjustments, samples taking different AVLTs were assumed to be randomly equivalent (see Limitations), such that verbal learning ability estimates were placed on the same scale using a ‘random equivalent groups’ linking design. The relative difficulty of all items across tests was inferred on a single ability scale, and tables of equivalent raw scores were linked across AVLTs.

### Transparency and Openness

Raw data are available upon reasonable request pending study approval and data transfer agreements between participating institutions. Code used for analysis and online tool creation are available online.

## Acknowledgements

· Spanish Ministry of Science and Innovation, Instituto de Salud Carlos III
  ○ PI15-00852
  ○ PI18-00945
  ○ JR19-00024
  ○ PI17-00481
  ○ PI20-00721
  ○ Sara Borrell contract (CD19-00149)
· European Union
  ○ NextGenerationEU (PMP21/00051)
  ○ PI19/01024
  ○ Structural Funds
  ○ Seventh Framework Program
  ○ H2020 Program under the Innovative Medicines Initiative 2 Joint Undertaking: Project PRISM-2 (Grant agreement No.101034377)
  ○ Project AIMS-2-TRIALS (Grant agreement No 777394)
  ○ Horizon Europe
· National Institutes of Health
  ○ U01MH124639
  ○ P50MH115846
  ○ R01MH113827
  ○ R25MH080663
  ○ K08MH068540
  ○ R01NS100973
  ○ R01EB006841
  ○ P20GM103472
  ○ RO1MH083553
  ○ T32MH019535
  ○ R01 HD061504
  ○ RO1MH083553
  ○ R01AG050595
  ○ R01AG076838
  ○ R01AG060470
  ○ R01AG064955
  ○ P01AG055367
  ○ K23MH095661
  ○ R01MH094524
  ○ R01MH121246
  ○ T32MH019535
  ○ R01NS124585
  ○ R01NS122827
  ○ R61NS120249
  ○ R01NS122184
  ○ U54EB020403
  ○ R01MH116147
  ○ R56AG058854
  ○ P41EB015922
  ○ R01MH111671
  ○ P41RR14075
  ○ M01RR01066
  ○ R01EB006841
  ○ R01EB005846
  ○ R01 EB000840
  ○ RC1MH089257
  ○ U24 RR021992
  ○ NCRR 5 month-RR001066 (MGH General Clinical Research Center)
· NSF
  ○ 2112455
· Madrid Regional Government:
  ○ B2017/BMD-3740 AGES-CM-2
· Dalhousie Medical Research Foundation
· Research Nova Scotia
  ○ RNS-NHIG-2021-1931
· US Department of Defense:
  ○ Award #AZ150145
· US Department of Veterans Affairs
  ○ 1I01RX003444
· NJ Commission on TBI Research Grants
  ○ CBIR11PJT020
  ○ CBIR13IRG026
· Department of Psychology, University of Oslo
· Sunnaas Rehabilitation Hospital
  ○ HF F32NS119285
· Canadian Institutes of Health Research:
  ○ Grant 166098
· Neurological Foundation of New Zealand
· Canterbury Medical Research Foundation, University of Otago.
· Biogen US
  ○ Investigator-initiated grant
· Italian Ministry of Health:
  ○ RF-2019-12370182
  ○ Ricerca Corrente RC 23
· National Institute on Aging:
· National Health and Medical Research Council
  ○ Investigator Grant APP1176426
· PA Health Research:
  ○ Grant SAP #4100077082 to Dr. Hillary
· La Caixa Foundation
  ○ ID: 100010434, fellowship code: LCF/BQ/PR22/11920017
· Research Council of Norway
  ○ 248238
· Health Research Council of New Zealand
  ○ Sir Charles Hercus Early Career Development (17/039)
  ○ 14-440
· South-Eastern Norway Regional Health Authority
  ○ 2018076
· Norwegian ExtraFoundation for Health and Rehabilitation
  ○ 2015/FO5146
  ○ 2015044
· Stiftelsen K.G. Jebsen
  ○ SKGJ MED-02
· German Research Foundation:
  ○ DFG grant FOR2107 to Andreas Jansen, JA 1890/7-1, JA 1890/7-2
  ○ DFG grant FOR2107 to Igor Nenadić, NE2254/1-2,NE2254/3-1,NE2254/4-1
  ○ DFG grant FOR2107, KI588/14-1 and FOR2107, KI588/14-2
  ○ DFG, grant FOR2107 DA1151/5-1 and DA1151/5-2
  ○ SFB-TRR58, Projects C09 and Z02
· Central Norway Regional Health Authority (RHA) and the Norwegian University of Science and Technology (NTNU)
· National Health and Medical Research Council
  ○ APP1020526
· Brain Foundation
· Wicking Trust
· Collie Trust
· Sidney and Fiona Myer Family Foundation.
· U.S. Army Medical Research and Materiel Command (USAMRMC)
  ○ Award #13129004
· Department of Energy
  ○ DE-FG02-99ER62764
· Mind Research Network
· National Association for Research in Schizophrenia and Affective Disorders
  ○ Young Investigator Award
· Blowitz Ridgeway and Essel Foundations
· NOW ZonMw TOP 91211021
· UCLAS Easton Clinic for Brain Hellath
· UCLA Brain Injury Research Center
· Stan and Patty Silver
· Clinical and Translational Research Center
  ○ UL1RR033176
  ○ UL1TR000124
· Mount Sinai Institute for NeuroAIDS Disparities
· VA Rehab SPIRE
· CDMRP PRAP
· VA RR&D IK2RX002922
· Veski Fellowship
· Femino Foundation grant
· Fundación Familia Alonso
· Fundación Alicia Koplowitz
· CIBERSAM, Madrid Regional Government (B2017/BMD-3740 AGES-CM-2)
· 2019R1C1C1002457, 21-BR-03-01
· 2020M3E5D9079910, 21-BR-03-01
· Interdisciplinary Center for Clinical Research (IZKF) of the medical faculty of Münster

## Author Contributions

### Designed research

Eamonn Kennedy, Hannah Lindsey, Pui-Wa Lei, Paul M Thompson, David F Tate, Frank G Hillary, Emily L Dennis, Elisabeth A Wilde

### Performed research

Eamonn Kennedy, Shashank Vadlamani, Hannah M Lindsey, Emily L Dennis, Elisabeth A Wilde

### Contributed data

Maheen Adamson, Martin Alda, Silvia Alonso-Lana, Sonia Ambrogi, Tim J Anderson, Celso Arango, Robert F Asarnow, Mihai Avram, Rosa Ayesa-Arriola, Talin Babikian, Nerisa Banaj, Laura J Bird, Stefan Borgwardt, Amy Brodtmann, Katharina Brosch, Karen Caeyenberghs, Vince D Calhoun, Nancy D Chiaravalloti, David X Cifu, Benedicto Crespo-Facorro, John C Dalrymple-Alford, Kristen Dams-O’Connor, Udo Dannlowski, David Darby, Nicholas Davenport, John DeLuca, Covadonga M Diaz-Caneja, Seth G Disner, Ekaterina Dobryakova, Stefan Ehrlich, Carrie Esopenko, Fabio Ferrarelli, Lea E Frank, Carol Franz, Paola Fuentes-Claramonte, Helen Genova, Christopher C Giza, Janik Goltermann, Dominik Grotegerd, Marius Gruber, Alfonso Gutierrez-Zotes, Minji Ha, Jan Haavik, Charles Hinkin, Kristen R Hoskinson, Daniela Hubl, Andrei Irimia, Andreas Jansen, Michael Kaess, Xiaojian Kang, Kimbra Kenney, Barbora Keřková, Mohamed Salah Khlif, Minah Kim, Jochen Kindler, Tilo Kircher, Karolina Knížková, Knut K Kolskår, Denise Krch, William S Kremen, Taylor Kuhn, Veena Kumari, Jun Soo Kwon, Roberto Langella, Sarah Laskowitz, Jungha Lee, Jean Lengenfelder, Spencer W Liebel, Victoria Liou-Johnson, Sara M Lippa, Marianne Løvstad, Astri Lundervold, Cassandra Marotta, Craig A Marquardt, Paulo Mattos, Ahmad Mayeli, Carrie R McDonald, Susanne Meinert, Tracy R Melzer, Jessica Merchán-Naranjo, Chantal Michel, Rajendra A Morey, Benson Mwangi, Daniel J Myall, Igor Nenadić, Mary R Newsome, Abraham Nunes, Terence O’Brien, Viola Oertel, John Ollinger, Alexander Olsen, Victor Ortiz García de la Foz, Mustafa Ozmen, Heath Pardoe, Marise Parent, Fabrizio Piras, Federica Piras, Edith Pomarol-Clotet, Jonathan Repple, Geneviève Richard, Jonathan Rodriguez, Mabel Rodriguez, Kelly Rootes-Murdy, Jared Rowland, Nicholas P Ryan, Raymond Salvador, Anne-Marthe Sanders, Andre Schmidt, Jair C Soares, Gianfranco Spalleta, Filip Španiel, Alena Stasenko, Frederike Stein, Benjamin Straube, April Thames, Florian Thomas-Odenthal, Sophia I Thomopoulos, Erin Tone, Ivan Torres, Maya Troyanskaya, Jessica A Turner, Kristine M Ulrichsen, Guillermo Umpierrez, Elisabet Vilella, Lucy Vivash, William C Walker, Emilio Werden, Lars T Westlye, Krista Wild, Adrian Wroblewski, Mon-Ju Wu, Glenn R Wylie, Lakshmi N Yatham, Giovana B Zunta-Soares,

### Analyzed data

Eamonn Kennedy, Shashank Vadlamani, Pui-Wa Lei

### Wrote paper

Eamonn Kennedy

### Edited paper

All authors

