## Supplement for "Bridging Big Data: Procedures for Combining Non-equivalent Cognitive Measures from the ENIGMA Consortium"

**Supplementary Information for : Standards for Converting Scores across Common Auditory Verbal Learning Tasks: An ENIGMA Mega-Analysis.**

Table S1: Summary characteristics of the underlying datasets (citations) (C = Control, LC = LIMBIC-CENC) Full Cohort descriptions are available in Supplementary Table S1. NDA datasets refer to multiple individual studies from the NIMH Data Archive. 'Mean Total Raw Score' refers to the raw sum of trials over the number of exposures in the learning portion of the tests. *No individual reference because data come from multiple studies.

| Dataset | Country | Lang. | Clinical Population (C=Control) | N | Female (%) | Age median (range) | Measure | Mean TrialsTot. Raw Score (SD) |
|---|---|---|---|---|---|---|---|---|
| 1. VETSA(*1*) | USA | English | TBI | 1291 | 0 | 55 (51-61) | CVLT | 42.7 (9.6) |
| 2. NDA Psych | USA | English | C | 1051 | 53 | 29 (18-61) | CVLT | 42.5 (9.6) |
| 3. NICoE(*2*) | USA | English | C, TBI | 1039 | 2 | 39 (20-59) | CVLT | 45.0 (9.8) |
| 4. LC Richmond(*3*) | USA | English | C, TBI | 343 | 18 | 45 (24-71) | CVLT | 43.5 (9.8) |
| 5. STROKE MRI(*4*) | Norway | Norwegian | C | 333 | 62 | 65 (20-94) | CVLT | 43.0 (9.8) |
| 6. LC Tampa(*3*) | USA | English | C, TBI | 284 | 15 | 37 (22-62) | CVLT | 44.0 (10.0) |
| 7. LC Baylor(*3*) | USA | English | C, TBI | 241 | 9 | 33 (22-54) | CVLT | 43.9 (10.0) |
| 8.SATURN/DEFEND(*5*) | USA | English | C, TBI | 233 | 6 | 31 (22-62) | CVLT | 42.8 (9.7) |
| 9. LC San Ant.(*3*) | USA | English | C, TBI | 218 | 12 | 39 (22-70) | CVLT | 44.1 (9.6) |
| 10. iSCORE(*6*) | USA | English | TBI | 209 | 11 | 34 (19-59) | CVLT | 43.8 (9.4) |
| 11.TRACK_UCSF(*7*) | USA | English | TBI | 195 | 32 | 36 (18-80) | CVLT | 42.3 (9.9) |
| 12. LC Portland(*3*) | USA | English | C, TBI | 156 | 13 | 38 (24-72) | CVLT | 43.6 (10.2) |
| 13. LC Fort Belv.(*3*) | USA | English | C, TBI | 124 | 11 | 41 (24-58) | CVLT | 44.8 (9.4) |
| 14. COBRIT(*8*) | USA | English | TBI | 112 | 62 | 31 (18-64) | CVLT | 43.4 (9.8) |
| 15. Inst. Pere M.(*9*) | Spain | Spanish | C | 93 | 100 | 40 (18-61) | CVLT | 42.9 (9.7) |
| 16. TRACK Pitt(*7*) | USA | English | TBI | 81 | 28 | 50 (18-79) | CVLT | 39.4 (9.5) |
| 17. BergenADHD(*10*) | Norway | Norwegian | C | 74 | 58 | 28 (19-45) | CVLT | 41.9 (9.9) |
| 18. GSU(*11*) | USA | English | C | 71 | 87 | 66 (60-80) | CVLT | 40.3 (9.5) |
| 19. SNUH(*12*) | S. Korea | Korean | C | 71 | 37 | 22 (18-48) | CVLT | 43.9 (9.9) |
| 20. MPLS VA(*5*) | USA | English | C, TBI | 68 | 10 | 38 (24-71) | CVLT | 42.7 (10.2) |
| 21. mSMTDODTBI(*13*) | USA | English | C, TBI | 64 | 28 | 43 (20-65) | CVLT | 42.9 (9.6) |
| 22. Dal.Nunes(*14*) | Canada | English | C | 61 | 62 | 24 (17-65) | CVLT | 43.0 (9.8) |
| 23. UBC(*15*) | Canada | English | C | 60 | 50 | 22 (16-37) | CVLT | 43.5 (9.7) |
| 24. LETBI(*16*) | USA | English | TBI | 48 | 58 | 48 (21-74) | CVLT | 43.7 (13.9) |
| 25. UCSD-EPI(*17*) | USA | English | C | 45 | 62 | 31 (19-72) | CVLT | 44.9 (9.5) |
| 26. FIDMAG(*9*) | Spain | Spanish | C | 36 | 64 | 37 (19-53) | CVLT | 46.3 (10.1) |
| 27. TRACK UMC(*7*) | USA | English | TBI | 20 | 15 | 47 (18-78) | CVLT | 42.6 (10.1) |

| | | | | | | | | |
|---|---|---|---|---|---|---|---|---|
| 28. BaselCHR(*18*) | Switz. | German | C | 13 | 69 | 27 (23-31) | CVLT | 49.0 (10.1) |
| 29. UCLA-HEPC(*19*) | USA | English | C | 185 | 22 | 56 (25-73) | HVLT | 22.3 (4.2) |
| 30. CARE(*20*) | USA | English | TBI | 156 | 37 | 20 (17-23) | HVLT | 23.1 (4.1) |
| 31. FURT(*21*) | Germany | German | C | 131 | 53 | 36 (20-65) | HVLT | 24.0 (4.0) |
| 32. Brunel(*22*) | UK | English | C, TBI | 111 | 0 | 24 (18-65) | HVLT | 23.3 (4.2) |
| 33. PSU(*23–25*) | USA | English | TBI | 93 | 37 | 60 (18-84) | HVLT | 23.5 (4.7) |
| 34. COBRE(*26*) | USA | English | C | 84 | 27 | 38 (18-65) | HVLT | 23.4 (4.1) |
| 35. IISGM* | Spain | Spanish | C | 73 | 44 | 21 (16-39) | HVLT | 23.3 (4.1) |
| 36. NDA SZ | USA | English | C | 66 | 59 | 56 (18-75) | HVLT | 23.9 (4.5) |
| 37. UCLA-NNFC(*27*) | USA | English | C | 62 | 42 | 50 (21-76) | HVLT | 23.0 (4.0) |
| 38. RAP(*28*) | USA | English | C | 58 | 40 | 22 (16-36) | HVLT | 22.7 (4.1) |
| 39. MCIC UNM(*29*) | USA | English | C | 41 | 22 | 23 (18-60) | HVLT | 23.3 (4.2) |
| 40. CANVAS(*30*) | Australia | English | C | 39 | 38 | 69 (30-89) | HVLT | 25.4 (4.5) |
| 41. MCIC UMN(*29*) | USA | English | C | 24 | 42 | 29 (18-60) | HVLT | 24.1 (4.3) |
| 42. MCIC MGH(*29*) | USA | English | C | 20 | 40 | 41 (20-57) | HVLT | 24.3 (4.2) |
| 43. HCP aging(*31*) | USA | English | C | 701 | 56 | 58 (36-90) | RAVLT | 41.8 (8.7) |
| 44. FOR MR(*32*) | Germany | German | C | 571 | 63 | 32 (18-69) | RAVLT | 44.2 (8.5) |
| 45. FOR MS(*32*) | Germany | German | C | 387 | 66 | 26 (18-66) | RAVLT | 44.8 (8.4) |
| 46. ADNIDoD(*33*) | USA | English | C, TBI | 289 | 1 | 69 (60-87) | RAVLT | 42.2 (8.4) |
| 47. NPL FSL(*34*) | Italy | Italian | C | 239 | 50 | 59 (16-87) | RAVLT | 41.7 (8.5) |
| 48. PAFIP(*35*) | Spain | Spanish | C | 201 | 39 | 29 (16-60) | RAVLT | 43.7 (8.6) |
| 49. IDOR | Brazil | Portug. | C, TBI | 191 | 64 | 68 (42-91) | RAVLT | 40.9 (8.3) |
| 50. NIMH CZ(*36*) | Czechia | Czech | C | 106 | 63 | 28 (18-54) | RAVLT | 45.0 (8.2) |
| 51. erTMS(*37*, *38*) | USA | English | TBI | 33 | 15 | 47 (26-69) | RAVLT | 43.6 (8.6) |
| 52. FETZ Bern(*39*) | Switz. | German | C | 6 | 17 | 18 (16-40) | RAVLT | 47.1 (8.5) |
| 53. MCIS(*40*) | Australia | English | C | 4 | 25 | 72 (58-79) | RAVLT | 38.3 (9.8) |

Table S2: Inclusion/Exclusion Criteria for each sub-study.

| Study | Description | Inclusion Criteria | Exclusion Criteria |
|---|---|---|---|
| ***ADNIDoD*** | The Alzheimer's Disease Neuroimaging Initiative, Department of Defense. | All subjects must be Veterans of the Vietnam War, 50-90 years of age, must live within 150 miles of the closest ADNI clinic in subject's area. TBI Subjects must have a documented history of moderate-severe non-penetrating TBI, which occurred during military service in Vietnam (identified from the Department of Defense or VA records). | All: MCI/Dementia, presence of PTSD by SCID-I for DSM-IV-TR criteria, or a CAPS score of >30 (Both current and/or a history of PTSD will be excluded). Control: Documented or self-report history of TBI, history of PTSD, MRI-related exclusions. |
| ***BaselCHR*** | Basel Study of Brain Dysfunctions in Subjects at Clinically High-risk for Psychosis. | For CHR: DSM-III-R criteria; for FEP: ICD-10/DSM-IV. | History of previous psychotic disorder, psychotic symptoms secondary to an organic disorder, substance abuse (except nicotine), psychotic symptoms associated with an affective psychosis or a borderline personality disorder, age younger than 18 years, inadequate knowledge of the German language, and IQ less than 70 as measured by the Mehrfachwahl Wortschatz Test Form B. |
| ***Bergen ADHD*** | Study of difficulties related to inhibition and set-shifting in adults with ADHD + a study investigating verbal memory function with relation to working memory (WM) and response inhibition (RI) in adults with ADHD. | The participants with ADHD (at least 18 years old) were recruited as part of a study that included participants from a national registry of adults diagnosed in Norway from 1997 to May 2005. Diagnostic assessment was conducted by three national expert committees for ADHD/hyperkinetic disorder. Clinicians with specialized experience in diagnosing ADHD served on the committees. The procedure for the referral required patient records with thorough descriptions of current symptoms and functioning. The committees reviewed diagnosis of ADHD. | There were no formal exclusion criteria. |
| ***Brunel*** | Brunel University Study of Schizophrenia including health controls | Schizophrenia: Adults (18-65 years) with a DSM-IV diagnosis of schizophrenia or schizoaffective disorder in a chronic illness phase (not within 2 years of illness onset). Personality Disorder: Adults with a DSM-IV diagnosis of antisocial personality disorder and/or ICD-10 diagnosis of Dissocial Personality Disorder. Anxiety and depression: Adults with ICD-10 diagnosis. | Healthy Controls: Any neuropsychiatric conditions. Schizophrenia: Current substance use disorder; any neurological illness or severe head trauma; not stable on antipsychotic medication for at least 3 months. Personality Disorder: Current substance use disorder (past history of substance abuse permitted and present for many); any neurological illness or severe head trauma. |
| ***CARE*** | The NCAA?DoD Concussion Assessment, Research and Education Consortium | Highschool and collegiate football athletes. | 1) Any contraindication or injury that would prevent participation in the study protocol; 2) current psychotic disorder or narcotic use; 3) history or suspicion of a clinical condition known to be associated with cognitive impairments (e.g. epilepsy, moderate-to-severe TBI). |
| ***COBRE*** | The Center for Biomedical Research Excellence study of schizophreniza including health controls | Schizophrenia: adults (18-65 years) with a DSM-IV diagnosis of schizophrenia. Diagnostic information was collected using the Structured Clinical Interview used for DSM Disorders (SCID). Healthy volunteers: completed the SCID non-patient edition to rule out axis I conditions. | history of neurological disorder, history of mental retardation, history of severe head trauma, with more than 5 minutes loss of consciousness, history of substance abuse or dependence, within the last 12 months. |

| | | | |
|---|---|---|---|
| ***COBRIT*** | The citicoline brain injury treatment trial | 1) Non-penetrating traumatic brain injury; 2) Age 18-70; 3) English speaking; 4) able to swallow oral medication; 5) GCS 3-12 with GCS motor less than or equal to 5 or motor score = 6 with >10mm diameter intraparenchymal hemorrhage, acute hematoma thickness >5mm, subarachnoid hemorrhage, intraventricular hemorrhage, midline shift > 5mm, or GCS 13-15 meeting above CT criteria. | 1) Intubation with GCS motor = 6 and no CT criteria; 2) bilaterally fixed and dilated pupils; 3) pregnant or breastfeeding; 4) disease that interferes with outcome; 5) acetylcholinesterase inhibitor use; 6) citicoline use; 7) prisoners. |
| ***Dal.Nunes*** | Dalhousie University Study of structural MRI to identify bipolar disorders including health controls | BD 1 and BD 2, diagnosed according to DSM-IV. | Current mood episode, organic mood disorder/history of brain injury, inability to speak English. |
| ***LC, various sites*** | PTSD and TBI Neuroimaging Lab Study of Adult OEF/OIF Veterans | All: 18-65, OEF/OIF veterans, fluent in English, free of implanted metal objects or metal shards in eyes, antidepressant, sleep, and anti-anxiety medication permitted. | All: Axis I other than PTSD or MDD, current substance abuse or lifetime substance dependence (other than nicotine), high risk for suicide, claustrophobia, neurological disorders, learning disability or developmental delay, major medical conditions. |
| ***FETZ Bern*** | The Bern Early Recognition and Intervention Centre for mental crisis (FETZ Bern)?An 8-year evaluation | Adolescents and adults (FETZ Bern) aged from eight to 40 years with a population catchment area of 1.035 million in Bern, Switzerland. Routine demographic, diagnostic and service usage data were collected upon admission to the service. | Exclusion criteria are (i) past clinical diagnosis of any psychotic disorder according to DSM and ICD (ii) diagnosis of delirium, dementia, amnestic or other neurological disorders, and (iii) CNS affecting conditions. |
| ***FIDMAG/Inst. Pere Mata*** | Sisters Hospitallers Research Foundation Neuroimaging Study | Bipolar disorder: right-handed adults (18-60 years) with a DSM-IV diagnosis of Bipolar Disorder, euthymic during the last 3 months (HDRS-21=<8 and YMRS=<6). At least two previous affective episodes. Healthy volunteers: right-handed adults (18-60 years) with no present or past psychiatric diagnosis (according to the Structured Clinical Interview for DSM-IV). | History of drug or alcohol abuse/dependence during the last year (excluding nicotine), severe neurological disorder (e.g. epilepsy, multiple sclerosis), history of head trauma, medical conditions affecting cognition. For healthy volunteers, present or past psychiatric diagnosis or treatment with psychotropic medication (except for mild night sedation), or a diagnosis of major mental disorder in a first degree relative were also exclusion criteria. |
| ***FURT*** | Frankfurt Memory Study | Bipolar disorder; right-handed adults (18-60 years) with a DSM-V diagnosis of Bipolar Disorder. Euthymic during the last 3 months (MADRS=<7). Healthy voluenteers: right-handed adults (18-60 years) with no present or past psychiatric diagnosis (according to the SCID). | History of drug or alcohol abuse/dependence, severe neurological disorder, history of head trauma, medical conditions affecting cognition. For healthy volunteers, present or past psychiatric diagnosis or a diagnosis of major mental disorder in a first degree relative were also exclusion criteria. |
| ***GSU*** | Georgia State University | African American adults with a diagnosis of type 2 diabetes; 60 years of age or older. | 1) history of head injury or major neurological disorder, 2) current clinically significant levels of depression or anxiety, or history of other major psychiatric disorder, 3) significant and uncorrected sensory deficits (e.g., vision, hearing), and 3) current or previous history of substance abuse including alcohol. Female participants recruited for the study were not on estrogen replacement therapy. |
| ***HCP Aging*** | Human Connectome Project, Aging. Inclusion and Exclusion Criteria not provided. Literature indicates a “screen was performed for all potential participants to rule out major exclusionary health conditions, and diagnostic and service usage data were collected upon admission to the service.” | | |

| | | | |
|---|---|---|---|
| ***PAFIP*** | Comparative Study of Aripiprazole, Quetiapine and Ziprasidone in Treatment of First Episode Psychosis: 3-year Follow-up | The patients met the following criteria: (1) 15-60 years of age; (2) lived within the catchment area; (3) were experiencing their first episode of psychosis; (4) had no prior treatment with antipsychotic medication or, if previously treated, a total life-time of adequate antipsychotic treatment of less than 6 weeks; and (5) met the DSM?IV criteria for brief psychotic disorder, schizophreniform disorder, schizophrenia, or not otherwise specified (NOS) psychosis. The diagnoses were confirmed using the Structured Clinical Interview for DSM-IV (SCID-I) carried out by an experienced psychiatrist 6 months after the baseline visit. | Patients were excluded for any of the following reasons: (1) meeting DSM-IV criteria for drug dependence; (2) meeting DSM-IV criteria for mental disability; and (3) having a history of neurologic disease or head injury. |
| ***IDOR*** | D'Or Institute for Research and Education | Elderly patients were referred to clinical and neuropsychological evaluation at a specialized center, because of memory complaints (self-report and/or collateral report). Patients were sorted into 3 categories: normal aging, Mild Cognitive Impairment and Dementia after clinical and neuropsychological evaluation, using DSM-5 criteria. | Psychiatric or neurological disorders. |
| ***IISGM*** | IPSMaranon | All participants: (1) 7-40 years old; (2) Written informed consent. First Episode Psychosis and Clinical Risk groups had additional inclusion criteria, but were not used in this study. | All participants: (1) Pregnancy; (2) History of head injury resulting in unconsciousness; (3) Intellectual disability; (4) Pervasive developmental disorder or autism spectrum disorder; (5) Neurological disorder. Substance use or abuse is not an exclusion criterion. |
| ***iSCORE*** | Imaging Support for the Study of Cognitive Rehabilitation Effectiveness | US patients with objective neuroimaging abnormalities predicting cognitive rehabilitation therapy effectiveness. | Any participants with conditions preventing MRI procedures (i.e. claustrophobia, shrapnel, pregnancy), neurologic conditions (e.g., seizures, psychosis, etc.), history of TBI exceeding mild severity or spinal cord injury, or current narcotic medicine use were excluded from the study. |
| ***LETBI*** | Late Effects of TBI project | At least one year following most recent complicated mild, moderate or severe TBI. Additional funding in 2020 expands to include individuals with substantial lifetime exposure to multiple mTBI and/or RHI (recruitment stalled secondary to pandemic); RHI sample therefore small. | - |
| ***LC/LIMBIC-CENC*** | The Long-Term Impact of Military-Relevant Brain Injury Consortium, multiple sites | Age 18 or over. Prior military combat deployment with any level of combat exposure. | (1) a history of moderate or severe TBI as defined as initial Glasgow Coma Scale < 13, coma duration > « hr, post-traumatic amnesia duration > 24 hr, or traumatic intracranial lesion on head CT.<br>(2) a history of major neurologic or psychiatric disorder such as stroke, spinal cord injury, or schizophrenia; (Major will be defined as resulting in a significant decrement in functional status or loss of independent living capacity.) |

| | | | |
|---|---|---|---|
| ***FOR-MS*** | Marburg-Munster Affective Disorders Cohort Study | Study participants were aged between 18 and 65.<br>Patients: psychiatric lifetime diagnosis as confirmed using the Structural Clinical Interview for DSM-IV-TR (SCID-I).<br>Healthy controls: no history of mental or neurological diseases | Exclusion criteria comprised the presence of any neurological abnormalities, history of seizures, head trauma or unconsciousness, severe physical impairment (e.g. cancer, unstable diabetes, epilepsy etc.), pregnancy, hypothyroidism without adequate medication, claustrophobia, color blindness, and general MRI contraindications (e.g. metallic objects in the body). Further, lifetime diagnoses of alcohol dependence posed reason for exclusion. |
| ***FOR-MR*** | Marburg-Munster Affective Disorders Cohort Study | Study participants were aged between 18 and 65.<br>Patients: psychiatric lifetime diagnosis as confirmed using the Structural Clinical Interview for DSM-IV-TR (SCID-I).<br>Healthy controls: no history of mental or neurological diseases. | Exclusion criteria comprised the presence of any neurological abnormalities, history of seizures, head trauma or unconsciousness, severe physical impairment (e.g. cancer, unstable diabetes, epilepsy etc.), pregnancy, hypothyroidism without adequate medication, claustrophobia, color blindness, and general MRI contraindications (e.g. metallic objects in the body). Further, lifetime diagnoses of alcohol dependence posed reason for exclusion. |
| ***MCIC*** | Mind Clinical Imaging Consortium of schizophrenia | All subjects were between the ages of 18 and 60 and spoke English as their native language. To be included in the schizophrenia cohort, patients had to meet DSMIV diagnostic criteria for schizophrenia, schizoaffective disorder or schizophreniform disorder. Concerted effort was made to recruit patients early in the course of their illness and especially those who were antipsychotic drug na‹ve.<br>The healthy control subjects with no current or past history of psychiatric illness including substance abuse or dependence were matched within site to the patient cohort for age, sex, and parental education. | Both patients and controls were excluded if they had 1)an IQ less than 70 based on a standardized IQ test, 2)history of a head injury resulting in prolonged loss of consciousness, neurosurgical procedure, neurological disease, history of skull fracture, severe or disabling medical conditions, or 3) a contraindication for MRI scanning such as pregnancy or implant.<br>Control subjects who met criteria for current or past history of substance abuse or dependence were excluded from the study. |
| ***MCIS*** | The Characterising Mild Cognitive Impairment using multimodal biomarkers study | Aged 30-85, at least 6 years education, Subjective and objective cognitive impairment consistent with MCI. | lifetime history of schizophrenia, schizoaffective disorder, or bipolar disorder, current major depressive episode or a past history of major depressive disorder with a high risk of relapse, current drug or alcohol abuse/dependence, history of alcohol abuse/dependence 6. Any significant disease or unstable medical condition that could affect cognitive testing, other neurological illness. A history of stable epilepsy is not an exclusion criterion. |
| ***CANVAS*** | Cognition and Neocortical Volume After Stroke | Adults with acute ischemic stroke in any vascular territory or etiology confirmed on clinical imaging. | pre-existing cognitive impairment (based on participant, primary care practitioner, and informant) or severe neurological or psychiatric disease; MRI exclusions; primary hemorrhagic stroke, TIA, or no clinically confirmed stroke; or were unlikely to survive 3 years due to severe medical illness. |
| ***NDA Psych/SZ*** | NDA NIMH Archive data, downloaded July 2022 | - | - |

| | | | |
|---|---|---|---|
| ***mSMT DOD TBI & Cognitive Reserve/ Neural Substrate studies*** | Cognitive Reserve: Cognitive Reserve in Traumatic Brain Injury<br><br>Neural Substrates: Neural Substrates of Facial Emotion Processing in Individuals with Brain Injury<br><br>mSMT DOD TBI: A Randomized Clinical trial Examining the Efficacy of the Group Administered Mofified Story Memory Technique (mSMT) in Person with TBI | Cognitive Reserve: aged 19-55, one year post injury, >18 years old at time of injury, no history of prior neurologic insult or disease (prior head injury, stroke, unstable or uncontrolled seizures, or brain tumor), free of open head injuries or focal injuries to the left hemisphere language areas, no significant psychiatric history (eg schizophrenia or bipolar disorder), no history of language learning disability, no history of inpatient alcohol or drug abuse treatment.<br><br>Neural Substrates: have a diagnosis of TBI (TBI) or no history of neurological injury or disease (HC), between 18-65, can read and speak English fluently<br><br>mSMT DOD TBI: ages 21-65, at least one-year post injury, primary language must be English, 20/60 minimum acuity in worst eye (Snell Eye Exam), Intact language comprehension (Montreal Cobgnitive Assessment / MoCA-BLIND cutoff score of 18), Deficits in New Learning and Memory (open-trial Selective Reminding Test). | Cognitive Reserve: MRI contraindications (aneurysm clip, implanted neural stimulator, implanted cardiac pacemaker or auto-defibrillator; cochlear implant (internal hearing aids); ocular foreign body (e.g. metal shavings), insulin pump or any pre-existing eye conditions; history of brain surgery, neurologic lesion, history of migraines; pregnant; left-handed .<br><br>Neural Substrates: left-handed; MRI contraindications: any non-MRI compatible metal in any part of body such as aneurysm clips, cardiac pacemaker, cochlear implants, metal fragments. History of welding, claustrophobia, or being pregnant; neurological disorder other than one TBI, epilepsy, multiple sclerosis; history of schizophrenia or bipolar disorder; currently taking steroids, benzodiazepines, neuroleptics .<br><br>mSMT DOD TBI: history of multiple sclerosis, seizures, or any other significant neurological history; unstable or uncontrolled seizures. Participants on steroids and /or benzodiazepines.<br>Participants were excluded if they have been hospitalized due to COVID-19. Any MRI contraindications. Individuals with an active diagnosis of Major Depressive Disorder, Schizophrenia, Bipolar Disorder I or II. Subjects whose vision is significantly impaired. |
| ***MPLS VA (two studies: SATURN, DEFEND*** | Two combined study cohorts:<br><br>SATURN "Study of Aftereffects of Trauma: Understanding Response in National Guard";<br><br>DEFEND "Essential Features of Neural Damage in Mild Traumatic Brain Injury" | 1) previously deployed US military veteran during OEF/OIF (Operations Enduring Freedom/Iraqi Freedom), 2) clinically significant posttraumatic stress symptomatology during screening (later confirmed by semi-structured CAPS diagnostic interview and diagnostic consensus), 3) history of likely mild traumatic brain injury due to blast exposure (later confirmed through neuropsychological consensus review of participant self-report using MN-BEST assessment tool), 3) a lack of clinically significant posttraumatic stress symptomatology and a lack of likely mild traumatic brain injury (later confirmed through CAPS and MN-BEST diagnostic consensus), 4) participant sampling of Minnesota National Guard members and outpatients at the Minneapolis Veterans Affairs Health Care System (VAHCS), 5) If currently receiving mental health services, inclusion required not altering any ongoing care. | 1) a neurological condition prior to deployment 2) current DSM-IV-TR-defined psychotic disorder, 3) current or past DSM-IV-TR-defined substance dependence other than alcohol, caffeine, or nicotine, 4) current DSM-IV-TR-defined substance abuse, 5) testing positive for elevated blood alcohol content the day of the study, 6) current or past formal diagnosis of attention-deficit/hyperactivity disorder, 7) significant risk of suicidal or homicidal behavior, 8) current or past unstable medical condition, 9) head injury from a source other than blast with LOC >10 minutes, post-traumatic amnesia greater than one hour, skull fracture, positive neuroradiological findings, or hospitalization for more than 24 hours, or 10) has been a frequent boxer or kickboxer. |
| ***NICoE*** | NICoE TBI Neuroimaging Core Project | 1. History or evaluation of head trauma or post-concussive symptoms; 2. Active duty or DEERS eligible individuals (to include non-active duty, military healthcare beneficiaries); 3. Adult between the ages of 18 and 60; 4. Males and non-pregnant/non-breastfeeding females (due to neuroimaging). | 1. Traumatic brain injury patients who are unable to consent themselves; 2. Actively enrolled in other randomized controlled treatment trials where this study would interfere; 3. History of prior severe neurologic or psychiatric condition, such as psychosis, stroke, multiple sclerosis, or spinal cord injury; 4. Pregnancy (by history and urine assay); 5. Breastfeeding. |
| ***NPL FSL*** | Foundation Santa Lucia | (i) Diagnosis of interest depending on the study protocol; (ii)Age >18yrs; (iii)suitability for MRI scanning; (iv)written informed consent | (i) history of alcohol or drug abuse in the two years before the assessment; (ii) lifetime drug dependence; (iii) traumatic head injury with loss of consciousness; (iv) past or present major medical illness or neurological disorders; (v) Intellectual disability; (vi) |

| | | | |
|---|---|---|---|
| | | | pervasive developmental disorder |
| ***NIMH CZ*** | Early Stage Schizophrenia Outcome - Neurocognition | The sample included healthy controls (HC) and individuals with an ICD-10 diagnosis of a first-episode psychotic disorder (F20, F23, and F25) (FEP). All participants were aged 18 or over and provided their written informed consent. | For all: 1) diagnosis of ADHD, dyslexia, dysgraphia, dyscalculia, or dysortographia, 2) history of brain trauma, and 3) signs of an invalid neuropsycholigical assessment according to examiner notes, e.g., neuropsychological tests administered within the past six months, studies psychology or works in the field. For HC: 1) diagnosis of a psychiatric disorder, and 2) has a first-degree relative with a diagnosis of an ICD-10 psychotic disorder. For FEP: 1) duration of illness > 2 years, and 2) psychiatric comorbidity according to M.I.N.I. (other than substance abuse) |
| ***StrokeMRI*** | Oslo Study of Stroke with MRI | 1) Healthy volunteers: recruited through advertisement in newspapers, social media and word-of-mouth. 2) Stroke patients: Patients admitted to the Stroke Unit at Oslo University Hospital and at Diakonhjemmet Hospital, Oslo, Norway during 2013?2016 were invited to participate. Stroke was defined as any form of strokes of ischemic or hemorrhagic etiology. We included patients in the chronic stage defined as a minimum of 6 months since hospital admission. | 1) Exclusion criteria included history of stroke, dementia, or other neurologic and psychiatric diseases, alcohol- and substance abuse, medications significantly affecting the nervous system and contraindications for MRI. 2) Exclusion criteria included transient ischemic attacks (TIA), MRI contraindications and other neurological diseases diagnosed prior to the stroke. |
| ***NZP3*** | New Zealand Parkinson's Progression Programme | 1) Parkinson's disease; 2) Healthy volunteers. | 1) Parkinson's - atypical parkinsonian disorder or other CNS disorder; previous history of other neurological conditions including moderate or severe head injury, stroke, learning disability or vascular dementia; and major medical illness in the previous 6 months. 2) Healthy volunteers - history of CNS disorder or any neurological conditions, including moderate or severe head injury, learning disability, and major medical illness in the previous 6 months. |
| ***erTMS*** | Efficacy of Repetitive Transcranial Magnetic Stimulation for Improvement of Memory in Older Adults with TBI | (1) Age 50-75 years, with a high school education (2) History of mild or moderate TBI. (3) Must be in the chronic stable phase of recovery (>6 months post injury) with residual cognitive difficulties. (5) If on a psychotropic medication regimen, that regimen will be stable for at least 4 weeks prior to entry. (6) Has an adequately stable condition. (7) Females agree to use acceptable methods of birth control. (8) Compliance and consent. | (1) Diagnosed with Dementia (2) Pregnant or lactating female. (3) Unable to be safely withdrawn, from medications that substantially increase the risk of seizures (4) Have a cardiac pacemaker or a cochlear implant, device, or metal in the brain (6) Have a mass lesion, cerebral infarct or other active CNS disease (7) Known current psychosis as determined by DSM-IV coding in chart (Axis I, psychotic disorder, schizophrenia) or a history of a non-mood psychotic disorder (8) Diagnosis of Bipolar Affective Disorder I. (9) Current amnesic disorders. (10) Current substance abuse (11) Prior history of seizures (12) Severe TBI or open head injury (13) TBI within last 6 months (14) Participation in another concurrent clinical trial (15) Patients with prior exposure to rTMS or ECT (16) Active current suicidal intent. |

| | | | |
|---|---|---|---|
| ***TBI rTMS*** | Repetitive Transcranial Magnetic Stimulation (rTMS) to Improve Cognition in TBI | Inclusion Criteria : 1) Only PTA will be used to differentiate between mild and moderate TBI. Our main recruitment resources (VA Polytrauma services) employ clinical interview and O-LOG We will include only those patients who are 1 SD or more below the mean score on Trail Making Test B (TMT B), using the published Heaton et al., (2004) norms, in keeping with the Common Data Elements. This sample includes Veterans with blast or non-blast deployment-related injuries. | Exclusion Criteria: 1) Pregnancy, lactation. 2) Unable to be safely withdraw from medications that increase the risk of seizures. 3) Have a cardiac pacemaker or a cochlear implant or device 4) Have a mass lesion, cerebral infarct or other active CNS disease. 6) Known current psychosis as determined by DSM-IV coding in chart or a history of a non-mood psychotic disorder. 7) Diagnosis of Bipolar Affective Disorder 8) Current amnesic disorders. 9) Current substance abuse 10) Prior history of seizures 11) severe TBI of open head injury 12) TBI within last two months or in acute stage 13) participation in another concurrent clinical trial 14) Patients with prior exposure to rTMS/ECT 15) Active current suicidal intent or plan. |
| ***RAP*** | Risk Assessment for Psychosis | Eligibility criteria in all groups included: (1) ages between 12 and 35 for CHR subjects and 12-40 for FEP and HC; (2) no lifetime history of head injury or neurological disorder resulting in loss of consciousness for more than 1 min, (3) being able to speak English fluently enough to participate in clinical assessments and study procedures; (4) able to travel to Western Psychiatric Hospital to participate in the study; (5) no pregnancy, (6) no history of drug or alcohol dependence in the past 6 months. CHR and FEP groups had different criteria, but were excluded from this study. | Exclusion criteria for HC participants are as follows: (1) Axis-1 psychiatric disorder diagnosis, assessed with the structured clinical interview for DSM disorders (SCID); (2) high-risk syndrome diagnosis; (3) first-degree relatives with a diagnosed psychotic disorder. FEP Patients were excluded for any of the following reasons: 1) had a psychotic illness with a temporal relation to a substance use disorder; 2) co-morbidity of DSM-IV psychoactive substance dependence within the past 6?months; 3) substance abuse (other than cannabis and/or alcohol) within the past month; or 4) a temporal relationship between illness onset and head injury. |
| ***PSU*** | Penn State University | All studies: moderate/severe TBI, defined as post-resuscitation GCS 3-12 or positive CT findings if GCS>12 | PA-DOH: 410007708: at least 55 years of age and at least 5 years post-injury. Given that chronic traumatic encephalopathy has been observed in individuals as young as 45 years of age, in order to guarantee sensitivity to brain changes and possible degeneration we included individuals at least 55 years of age with no upper age restriction. For NIH Tr000127 and NJCBI 0120090178, age ranges included 18-75 years. For all studies, Individuals with significant neuropsychiatric disturbance (i.e., schizophrenia, bipolar disorder), or substance abuse requiring inpatient rehabilitation, or neurological disorder other than TBI were excluded. |
| ***RAPBI*** | Recovery After Pediatric Brain Injury | 1) Non-penetrating moderate-severe TBI (msTBI) classified according to intake or post-resuscitation Glasgow Coma Scale score between 3 and 12 (TBI group only); 2) between 8-19 years of age; 3) right-handed; 4) normal visual acuity or vision corrected with contact lenses/eyeglasses; and 5) English skills sufficient to understand instructions and be familiar with common words (the neuropsychological tests used in this study presume competence in English). | 1) history of neurological illness, such as prior msTBI, brain tumor, or severe seizures; 2) motor deficits that prevent the participant from being examined in an MR scanner (e.g., spasms, movement disorder); 3) history of psychosis, ADHD, Tourette?s syndrome, learning disability, intellectual disability, autism, or substance abuse, as these conditions are associated with cognitive impairments that might overlap with those caused by msTBI; 4) MRI contraindication. |

| | | | |
|---|---|---|---|
| ***SNUH*** | Seoul National University Hospital | For first-episode psychosis (FEP), an individual aged 16 to 40 years who satisfied the diagnosis of schizophreniform disorder, schizophrenia or schizoaffective disorder when assessed using the Structured Clinical Interview for the Diagnostic and Statistical Manual of Mental Disorders, Fourth Edition, Axis I Disorders (SCID-I), and the duration of psychotic illness was less than 2 years. For OCD, diagnosis were made according to the Diagnostic and Statistical Manual of Mental Disorders, fourth edition (DSM-IV) criteria and patients were either drug naive or unmedated for more than 4 weeks before assessment. | Potential HC participants were excluded when they had any first- to third-degree biological relatives with a psychotic disorder. Common exclusion criteria included substance abuse or dependence (except nicotine), neurological disease or significant head trauma, medical illness that could accompany psychiatric symptoms, and intellectual disability (intelligence quotient [IQ] < 70). |
| ***TRACK-TBI*** | TRACK-TBI | 1. Age 18 - 65 years inclusive; 2. History or evidence of TBI, according to DoD-VA criteria; 3. Glasgow Coma Scale (GCS) 3 - 15 after resuscitation in the ED; 4. Head CT with evidence of trauma-related abnormality (except for isolated epidural hematoma (EDH)); 5. Ability to undergo MRI within 48 hours of injury; 6. Ability to obtain informed consent from participant or Legally Authorized Representative (LAR) within 6 hours of injury; 7. Fluency in English or Spanish; ; | 1. Unstable respiratory or hemodynamic status; 2. Evidence of penetrating brain injury; 3. Isolated EDH; 4. Systemic traumatic injury not related to TBI; 5. Evidence of serious infectious, pneumonia; 6. Acute ischemic heart disease; 7. History of syncope or hypotension; 8. Systolic blood pressure (SBP) < 90 mm Hg, Diastolic blood pressure (DBP)< 40 mm Hg for longer than 5 minutes; 9. History or evidence of active malignancy; 10. History of pre-existing neurologic disorder, or other disorder that may confound interpretation of MRI or neuropsychological results; 11. History of pre-existing disabling mental illness, such as major depression or schizophrenia; 12. History or evidence of chronic heart/renal failure; 13. Low likelihood of follow-up 14. Women who are pregnant or breast-feeding; 15. Prisoners or patients in custody; 15. Patients on psychiatric hold (e.g. 5150, 5250). |
| ***UBC*** | | For bipolar group: Inclusion Criteria: Patients met DSM criteria for Bipolar I disorder based on assessment with MINI; first manic episode within the last 3 months, with or without psychosis, pure or mixed mania, with or without comorbidity (e.g. substance use history); age 14 to 35 years.<br>For healthy controls: Inclusion: Healthy individuals comparable to patient group on demographics. | For Bipolar group: Exclusion Criteria: inability to take part in neuropsychological testing (e.g. poor English, clinically unstable), evidence of a previous manic episode diagnosed retrospectively on structured interview or via collateral; history of major medical or neurological illness underlying their manic symptoms. For healthy controls: Exclusion: also received MINI; personal or family history of major psychiatric disorder in first-degree relatives; major medical or neurological illness affecting cognition |
| ***UCLA-NNFC*** | | 18 years (range 23?75), reported English as their primary language, scored ò26 on the Mini-Mental Status Exam (MMSE; Folstein et al. 1975), self-identified as African-American or Caucasian, and were able to provide informed consent | Participants were excluded if they reported either current abuse/dependence of cocaine or amphetamines, past stimulant abuse/dependence, or current/past diagnosis of a psychotic-spectrum disorder (assessed by a modified version of the SCID; Spitzer et al. 1995). Recent illicit drug use was assayed via urine toxicology as well as a self-report drug screen (i.e., Brief Drug History Questionnaire). Participants were excluded if they screened positive for stimulants or hallucinogens. Participants with CNS confounds (e.g., HIV-associated opportunistic infections or neurosyphilis), Hepatitis C coinfection, (confirmed by serology), major head injury (loss of consciousness >30 min), or contraindication to the safe use of MRI, were also excluded. |

| | | | |
|---|---|---|---|
| ***UCLA-HEPC*** | UCLA Hepatitis C Study | Inclusion criteria were: (a) 18 years of age or older, (b) proficient at reading and writing in English, and (c) reading proficiency at or above the 6th grade level. | Exclusion criteria were: (a) cirrhosis/liver failure assessed via blood tests or liver biopsy with model for end-stage liver disease (MELD) score > 12; (b) current or past psychotic spectrum disorder; (c) current moderate or severe major depressive disorder; (d) history of learning disorder, neurologic disorder (e.g., seizure disorder, stroke), head injury with loss of consciousness equal to or greater than 30 min, or any neurologic disease; (e) concurrent hepatitis A or B infection; (f) diagnosis of HIV as assessed via seropositive HIV antibody testing; (g) recent illicit drug use as assessed via urine toxicology; and (h) contraindication for MRI. |
| ***UCSD-EPI*** | | All participants were between the ages of 18-65. A diagnosis of TLE was established by a board-certified neurologist with expertise in epileptology, based on video-EEG telemetry, seizure semiology, and neuroimaging. Participants were included if they met the following criteria: 1) age 18-65, 2) English-speaking, 3) estimated premorbid IQ ò 70), 4) had available at least one memory score and 5) if they had no evidence of large structural lesions or visible extra-hippocampal pathology on MRI. | Healthy controls were excluded if they 1) reported a history of neurological (including TBI) or psychiatric diagnoses, 2) had a estimated intellectual function of less than 70 (WTAR standard score) and 3) performed below 2.5 standard deviations below the mean of the CVLT II normative sample. |
| ***Houston*** | | Specific inclusion criteria for the BD subjects are: a) BD proband with diagnosis of BD I, based on DSM-IV criteria, b) ages 18-65 years old; c) BD proband at any current mood state at the time of the study; d) BD proband preferably off pharmacological treatment at the time of study, but if not feasible, being on antidepressants and mood stabilizers (including anticonvulsants, typical and atypical antipsychotics, and lithium will be allowed. The inclusion criteria for healthy controls are: a) being mentally healthy, defined as no lifetime or present history of any axis I psychiatric disorder including alcohol or substance abuse/dependence, based on DSM-IV criteria; b) ages 21-50 years old. | Exclusion criteria for health control subjects are: a) history of any axis I psychiatric disorder in first-degree relatives, b) having taken a prescribed psychotropic medication ever. Exclusion criteria for the BD sibling pairs: a) diagnosis of Schizoaffective Disorder or Schizophrenia. Alcohol and substance abuse/ dependence (if in remission) and anxiety disorders are allowed; b) regular dose of benzodiazepines within two weeks of study; c) presence of current psychotic symptoms; d) pregnancy e) ineligibility or inability to the study. current major medical problems that affect brain anatomy, neurochemistry, or function, b) history of any brain diseases or head injury with loss of consciousness for any period of time; c) family history of hereditary neurologic disorder; d) floating metallic objects in the body (exclusion for MR studies); e) pregnancy. |
| ***VETSA*** | Vietnam Era Twin Study of Aging | At baseline, Participants were randomly recruited from the Vietnam Era Twin Registry aka VETR (by registry definition, then, both brothers had been in the US military at some point between 1965 and 1975; not "VA" patients--commmunity dwelling). Participants had to be between 50 to 59 years old when recruited; both brothers needed to agree to participate. Followups did not require pairs due to effects of aging (e.g. deaths, disability) that might prevent one from participating. at VETSA 2 and 3 attrition replacements (randomly recruited from VETR) were added who were age matched to that wave. | For MRI component, participants needed to meet safety criteria. |

40. MCIS: Characterising Mild Cognitive Impairment using multimodal biomarkers. *Monash University*, (available at https://research.monash.edu/en/projects/mcis-characterising-mild-cognitive-impairment-using-multimodal-bi).
